# *Grin2a* Hypofunction Disrupts Hippocampal Network Oscillations and E/I Balance Contributing to Cognitive Deficits

**DOI:** 10.1101/2024.11.09.622809

**Authors:** Hassan Hosseini, Jean C. Rodríguez Díaz, Dominique L. Pritchett, Morgan Sheng, Kevin S. Jones

**Affiliations:** Department of Pharmacology, University of Michigan Medical School, Ann Arbor, MI, USA; Neuroscience Graduate Program, University of Michigan Ann Arbor, MI, USA; Biology Department, Howard University, Washington, DC, United States of America; Stanley Center for Psychiatric Research, Broad Institute of MIT and Harvard, Cambridge, MA, USA; Department of Brain and Cognitive Sciences, Massachusetts Institute of Technology, Cambridge, MA, USA

## Abstract

**Background:** N-methyl-D-aspartate (NMDA) receptors, particularly those containing the GluN2A subunit, play a critical role in hippocampal-dependent learning and memory. The GluN2A subunit, encoded by the *Grin2a* gene, is essential for maintaining cognitive function, including working memory. In this study, we explore how *Grin2a* mutations impair working memory and disrupt hippocampal network oscillations and E/I balance.

**Methods:** *Grin2a* mutant mice were assessed for spatial working memory deficits using the 8-arm radial maze. We utilized multi-electrode arrays and whole-cell patch-clamp electrophysiology to evaluate network oscillation and synaptic inputs to pyramidal cells in *ex vivo* hippocampal slices. We performed an immunohistochemical analysis of hippocampal slices to evaluate changes in the abundance of GABAergic neurons.

**Results:** Our results demonstrate that *Grin2a* deficiency impairs spatial working memory and disrupts coupling of theta-gamma oscillations in the hippocampus. Moreover, *Grin2a* mutants express an overabundance of parvalbumin-expressing (PV+) interneurons that integrate into hippocampal circuits and destabilize excitatory/inhibitory (E/I) input to CA1 pyramidal neurons.

**Conclusion:** This study highlights the critical role of GluN2A-containing NMDA receptors in maintaining hippocampal network synchrony. Impairments in network synchrony and E/I balance within the hippocampus may drive the cognitive deficits observed in *Grin2a*-related disorders such as schizophrenia, epilepsy, and intellectual disability.

---

N-methyl-D-aspartate receptors (NMDARs), especially those containing the GluN2A subunit, are integral to hippocampal function, contributing to learning, memory, and network synchronization (Traynelis et al. 2010). Dysregulation of NMDAR signaling has been implicated in several psychiatric disorders, and mutations to the GRIN2A gene– which encodes the GluN2A subunit–increases the risk of schizophrenia 20-fold (Harrison and Weinberger 2005; Moghaddam and Javitt 2012; Singh et al. 2022). Transgenic *Grin2a* mouse models exhibit behavioral and physiological abnormalities reminiscent of schizophrenia, including impaired spatial memory (Sakimura et al. 1995; Kannangara et al. 2015). While synaptic plasticity deficits are central to these abnormalities, emerging evidence highlights that GluN2A-containing NMDARs also regulate critical neurophysiological processes, including network oscillations and excitatory/inhibitory (E/I) balance, which are essential for cognitive function (Hanson et al. 2020). This study will explore how disruptions in these processes can lead to a non-linear gene-dose response, where even partial loss of GluN2A function produces severe cognitive and physiological deficits.

Mutations in *Grin2a* can disrupt the function of GluN2A-containing NMDARs, leading to impaired gamma-band oscillations (GBOs) and theta-gamma phase-amplitude coupling (PAC) (Bertocchi et al. 2021), which are crucial for memory and cognitive processing. GBOs originate from synchronized neural activity and are predominantly driven by PV+ interneurons (Cardin et al. 2009). GBOs are crucial for memory encoding and intensify with increasing cognitive load (Montgomery and Buzsáki 2007). Schizophrenia frequently features abnormally high resting-state GBOs, and several studies have identified these aberrations in affected individuals (McNally and McCarley 2016; Hirano et al. 2015; Kwon et al. 1999). Elevated hippocampal GBOs are particularly associated with cognitive deficits, including impaired working memory, in schizophrenia patients (Tregellas et al. 2014). NMDAR antagonists further exacerbate these symptoms and replicate key features of schizophrenia, such as heightened resting-state GBOs, in both human and rodent models (Pinault 2008; Mann and Mody 2010; Kittelberger et al. 2012).

Importantly, NMDAR antagonists not only elevate resting-state GBOs but also disrupt their coupling with other brain rhythms (Abad-Perez et al. 2023). In the hippocampus, the amplitude of GBOs is modulated by the phase of theta-band oscillations (TBOs), a process known as phase-amplitude coupling (PAC). This coupling is critical for organizing the temporal dynamics of hippocampal circuits (Lisman and Idiart 1995). Theta-gamma PAC enhances the encoding and retrieval of spatial information by synchronizing neural assemblies, thereby facilitating memory storage and processing (Canolty and Knight 2010). Moreover, hippocampal theta-gamma PAC is crucial for working memory in humans (Daume et al. 2024), and disruptions in theta-gamma PAC are associated with cognitive impairments observed in dementia (Goodman et al. 2018) and schizophrenia (Barr et al. 2017).

This study investigated how *Grin2a* dysfunction impacts spatial working memory and the associated GBO and theta-gamma PAC in *ex vivo* hippocampal sections. Immunohistochemical analysis of hippocampal tissue revealed that *Grin2a* ablation increases the abundance of PV+ interneurons and patch-clamp recordings demonstrated an increase in inhibitory currents, disrupting the E/I balance in CA1 pyramidal neurons. Together, these findings offer crucial insights into the role of GluN2A-containing NMDARs in hippocampal network dynamics and provide a pathophysiological framework for understanding cognitive impairments associated with *Grin2a* mutations in psychiatric disorders. To our knowledge, this is the first study to directly investigate how *Grin2a* dysfunction affects both GBOs and its relationship to other frequency bands, offering new insights into pathophysiological mechanisms of cognitive impairments linked to schizophrenia.

## Methods

### Animals

Mice were housed in the University of Michigan’s animal care facilities under controlled temperature and lighting conditions (12/12 h light/dark cycle), with *ad libitum* access to food and water. All animal procedures adhered to the University of Michigan’s Institutional Animal Care and Use Committee (IACUC) guidelines and complied with NIH Guidelines for Animal Use. Mice were used between 3 and 6 weeks of age for network recording and 12 and 14 weeks of age for behavior and patch clamp recording. *Grin2a* knockout (KO) mice (Riken B6;129S-*Grin2a*^tm1Nak^; RBRC02256) (Kadotani et al. 1996) were obtained and outcrossed to C57BL/6J mice (RRID: IMSR_JAX:000664) to *Grin2a+/-* for three generations. *Grin2a*+/+ mice used in this study were offspring of crosses between C57BL/6J mice or the *Grin2a*+/+ mice that resulted from interbreeding Grin2a +/- mice. *Grin2a*+/- mice were obtained from interbreeding of *Grin2a*+/- mice.

*Grin2a*-/- mice were obtained from interbreeding *Grin2a*-/- or *Grin2a*+/- mice. The genotypes of all *Grin2a* mutant mice (henceforth referred to as *Grin2a* mice) were confirmed via real-time PCR assay (Transnetyx, Cordova, TN). A subset of data acquired from WT mice used in Fig. 2C or Fig. S2A-C were previously published (Rodríguez Díaz et al. 2023). Experimenters were blinded to the genotype of all mice until analyses were completed.

### Radial Arm Maze Test

The 8-arm radial maze utilizes the natural foraging behavior of rodents to assess spatial working memory (Olton and Samuelson 1976). The maze comprises an octagonal central platform (35 cm in diameter) with eight evenly spaced arms (60 cm long, 12 cm wide) extending outward, each capable of containing a food reward. This configuration allows for the evaluation of the rodent’s ability to recall and distinguish between visited and unvisited arms, thereby measuring spatial memory and learning capacity.

### Radial Arm Maze Test Protocol

Before the training sessions, food was withheld for 12 hours. The body weight of each mouse was maintained at no less than 85% of baseline weight. During each training session, mice were allowed to explore the arms of the maze to retrieve sucrose pellets (Dustless Precision Pellets®, Sucrose, Product No. F0026) until all eight arms had been visited. The maze was situated in an experimental room, with each arm distinguished by a different color and supplemented with fixed external visual cues. Mice received 10 training sessions, and were then tested, once per day for 10 consecutive days. Each test trial was conducted in three distinct phases: Target Phase: The mouse is placed in the maze with four arms pseudorandomly obstructed, allowing it to enter the remaining unblocked arms. Retention Phase: A mouse is placed in a new cage for 60 seconds. Test Phase: The mouse is reintroduced to the maze in which all arms are now unobstructed, and only previously obstructed arms from the target phase are baited with sucrose pellets. An entry into an arm is defined as the placement of all four paws into the arm.

### Radial Arm Maze Error Analysis

Working memory performance was video recorded and manually analyzed offline for reference memory errors, and working memory errors. Reference memory errors were defined as entries into arms that were never baited, whereas working memory errors were defined as times when mice re-entered arms that they had already visited during the target phase. These metrics were systematically analyzed to evaluate spatial working memory performance, following the methodology described by (Schmitt et al. 2003). The apparatus was sanitized with a 70% ethanol solution between trials to eliminate trace odors and maintain a consistent odor between trials. All procedures were conducted during the mouse light period to control for potential circadian influences.

### Slice electrophysiology

#### Multielectrode array studies

Electrophysiological recordings followed previously published methods (Rodríguez Díaz et al. 2023). In brief, 59-electrode perforated microelectrode arrays (pMEAs; Multichannel Systems, Reutlingen, Germany) were used to capture neural activity from 300 µm horizontal hippocampal sections. Mice were anesthetized with isoflurane, perfused with ice-cold NMDG-HEPES aCSF, and brain sections were prepared using a vibrating microtome (Leica VT1200). After incubation in NMDG aCSF (33°C, 10–12 min) and transfer to HEPES-buffered aCSF, sections were mounted on pMEAs and superfused at 5–7 ml/min with oxygenated aCSF (29–31°C). Signals were digitized at 20 kHz using a MEA2100 headstage, with baseline recordings acquired for 1 hour.

Oscillatory activity was evoked by bath application of 400 nM kainate. The strategic placement of multiple electrodes within the CA1 and CA3 subfields enabled the investigation of interactions both within and between hippocampal subfields (Rodríguez Díaz et al. 2023). Horizontal hippocampus sections maintain longitudinal connectivity between CA3 and CA1 (Boehlen et al 2009). All solutions were maintained at pH 7.4 by saturation with a gas mixture of 95% O_2_ / 5% CO_2_. Recordings were digitized at 20 kHz using a MEA2100 headstage (Multichannel Systems, Reutlingen, Germany). Digitized signals were stored on a personal computer for offline analysis.

#### Patch-clamp studies

Whole-cell patch clamp recordings were obtained from CA1 pyramidal neurons in *ex vivo* coronal hippocampal sections. Borosilicate glass pipettes (Sutter Instrument 1B120F-4) with resistances ranging from 3–5 MΩ were fabricated using a laser micropipette puller (Sutter Instrument Model P-2000). The pipettes were filled with an intracellular solution, adjusted to a pH of 7.3 and 290 mOsm, composed of in mM: 120 Cs-methanesulfonate, 5 CsCl, 10 Na_2_-phosphocreatine, 10 HEPES, 4 MgATP, 0.3 GTP-Na, and 5 QX314. For a subset of cells, 0.3% biocytin was added to the intracellular solution to facilitate post-hoc visualization. Whole-cell patch clamp recordings were acquired using a MultiClamp 200B amplifier. Signals were filtered at 8 kHz and digitized at 20 kHz using a Digidata 1550A digitizer (Molecular Devices, San Jose, CA). Digitized signals were stored on a personal computer for offline analysis. Spontaneous excitatory postsynaptic currents (sEPSCs) and spontaneous inhibitory postsynaptic currents (sIPSCs) were recorded in aCSF (in mM: 124 NaCl, 2.5 KCl, 2 CaCl_2_ 2 MgCl_2_, 1.2 NaH_2_PO_4_, 5 HEPES, 24 NaHCO_3_, 12.5 dextrose) at holding potentials of −70 mV and 0 mV, respectively. Electrophysiological traces were filtered at 1 kHz. Templates for semi-automatic event detection were generated based on the mathematical mean of manually curated waveforms using Clampfit software. We quantified the net contribution from sEPSCs and sIPSCs by calculating average net charge transfer (mean area under the curve x mean frequency). This allowed us to plot the ratio of total charge transfer for both excitatory and inhibitory synaptic activity.

### Data processing and statistical analysis

#### Gamma and theta band power

Theta band oscillations (TBOs) between 4-10 Hz and “low” gamma band oscillations between 25 and 59 Hz were analyzed (Colgin et al. 2009). Frequencies from 25 to 59 Hz were particularly emphasized in our analysis to avoid interference from 60 Hz line noise. Absolute power within the gamma band was calculated by integrating the power spectrum over theta and low gamma frequency range. Data preprocessing was conducted on a workstation, with coherence and periodicity analyses of GBOs performed in MATLAB (MathWorks, Natick, MA) and subsequently exported to GraphPad Prism for statistical analysis and plotting, except where otherwise noted. Local field potentials were isolated from microelectrode array (MEA) recordings by applying a low-pass infinite impulse response (IIR) Butterworth filter at 300 Hz and down-sampled to 1 kHz on each channel.

Spectrograms were generated by convolving the signals with a Morlet wavelet as follows: 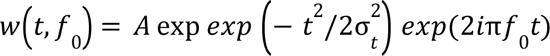, where A is a normalization factor equal to 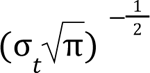. The width of the wavelet was set to 25, 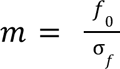 with 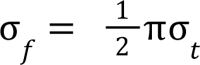. Power analysis was performed using multi-taper spectrum analysis with custom MATLAB scripts and multi-tapered Fourier estimation (mtspecgramc) (19)(Chronux Home, n.d.).

GBO onset was defined as the time required for GBO power to reach 90% of the maximum and was calculated using the rise time function as described above. GBO Coherence_90_ was fitted to the sigmoid function 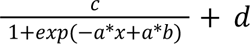. The quality factor value (Q value) of the oscillations (21) was calculated by the equation Q=f0/B, where f0 is the peak frequency and B is the bandwidth at 50% of maximum peak power.

#### Gamma Band Coherence

To calculate the power-power coherence of kainate induced GBOs, we used the cohgramc function frome Chronux toolbox (version 2.11), and calculated the autocorrelation on each channel, Sx(f) and Sy(f), and the cross-spectrum between the channels Sxy(f), which is calculated as follows: 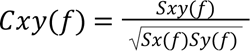 (20). Fold change in GBO coherence was calculated dividing the the GBO coherence during last two minutes of bath application of kainate by the GBO coherence during the last five minutes of baseline activity in vehicle:

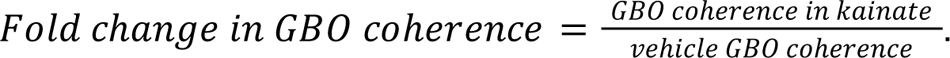

Interhemispheric, zero-phase-lag coherence of GBOs was computed between all MEA electrodes using a multi-tapered approach.

#### Phase-Amplitude Coupling (PAC) Analysis

Phase-amplitude coupling (PAC) was analyzed using a custom Python script, leveraging functions from the Tensorpac toolbox (https://etiennecmb.github.io/tensorpac/). We chose phase frequencies between 2 to 25 Hz in 0.2 Hz increments, and amplitude frequencies between 1 to 100 Hz in 1 Hz increments. Coupling strength was estimated using normalized direct PAC (ndPAC) method which builds on the mean vector length approach by incorporating a z-scored normalized amplitude and a statistical test based on a normal distribution of phase values from -π to π (Ozkurt, 2012). Our results report the coupling of low-frequency gamma oscillations (25–59 Hz) and theta phase (4–10 Hz). We then used Kruskal Wallis test to compare ndPAC strength before and after kainate application.

### Spike Sorting

Spike extraction from the local field potential (LFP) signal was performed using a customized function with specific input parameters. The data were bandpass filtered between 200 Hz and 10 kHz, and a sample window of 4.0 ms was defined to facilitate signal extraction. A threshold of 5x the average root mean square of signal was set to detect potential spikes, which were then aligned at the maximum amplitude. Spikes that exceeded the threshold were retained, others were excluded.

After spike extraction, waveforms were normalized and subjected to principal component analysis (PCA) for dimensionality reduction. Cluster analysis was conducted using the K-Means (k = 2) clustering algorithm, with the optimal number of clusters determined by the Bayesian Information Criterion. Neurons were classified into putative excitatory (pE) or putative inhibitory (pI) units based on their after-hyperpolarization characteristics of the mean waveform. Specifically, neurons exhibiting after-hyperpolarization amplitudes of less than 1.3 times their baseline level were classified as Ep, whereas those with higher values were classified as Ip (Sirota et al. 2008).

### Spike-Field Coherence

Spike-field coherence was calculated using the phase-locking value (PLV) technique, yielding values ranging from 0 (no coherence) to 1 (perfect coherence). LFP signals were bandpass-filtered in the theta frequency range (4-10 Hz), and the phase of the filtered signal was obtained through a Hilbert transform. The corresponding LFP phase was determined for each spike event and binned into 20 intervals and spike occurrences were normalized across phase bins to compute the PLV. To assess the significance of coherence values, we employed a bootstrapping method, generating 1,000 random spike-phase distributions for comparison with observed values. Inclusion criteria for slice analysis of phase-amplitude coupling (PAC) and spike sorting required the presence of at least three electrodes placed in both the CA1 and CA3 regions.All analyses were conducted using custom Python scripts incorporating libraries such as *scipy* and *matplotlib*, *seaborn*, and *sklearn*. Code available upon request.

### Immunohistochemistry

Hippocampal sections were fixed in 4% paraformaldehyde for 24 hours at 4°C, and transferred to phosphate-buffered saline (PBS) and cryoprotected in 30% sucrose until sectioning. Free-floating sections (40 µm thick) were prepared using a freezing microtome and blocked in 10% normal goat serum with 0.5% Triton X-100 in PBS for one hour at ambient temperature. Sections were incubated with an α-parvalbumin primary antibody (1:500, Millipore) overnight at 4°C. After washing, sections were incubated with fluorophore-conjugated secondary antibodies (Alexa Fluor 488, 1:1000, Thermo Fisher) for two hours at ambient temperature. Stained sections were mounted on slides, coverslipped with Vectashield mounting medium, and visualized using a confocal microscope.

## Results

### Grin2a mutant mice exhibit impaired spatial working memory

We assessed spatial working memory in *Grin2a* mutant mice using the 8-arm radial maze, where mice differentiate between visited and unvisited arms (Olton and Samuelson 1976). *Grin2a* mutant mice (henceforth referred to as *Grin2a* mice) exhibited difficulties tracking visited arms during the target phase, where four arms were randomly blocked (**Fig. 1A**). By the third session, WT mice completed the task without making reference errors, while *Grin2a* mice continued to make errors throughout all sessions (**Fig. 1B**). Initially, WT and *Grin2a* mice made similar working memory errors. However, WT performance continuously improved, and by the ninth session, WT mice completed the task without errors. In contrast, *Grin2a*-/- and *Grin2a*+/- mice showed no improvement and made significantly more errors than WT mice by the sixth and seventh sessions, respectively (**Fig. 1C**). These results demonstrate the importance of GluN2A-containing NMDARs in spatial working memory.

**Figure 1.**
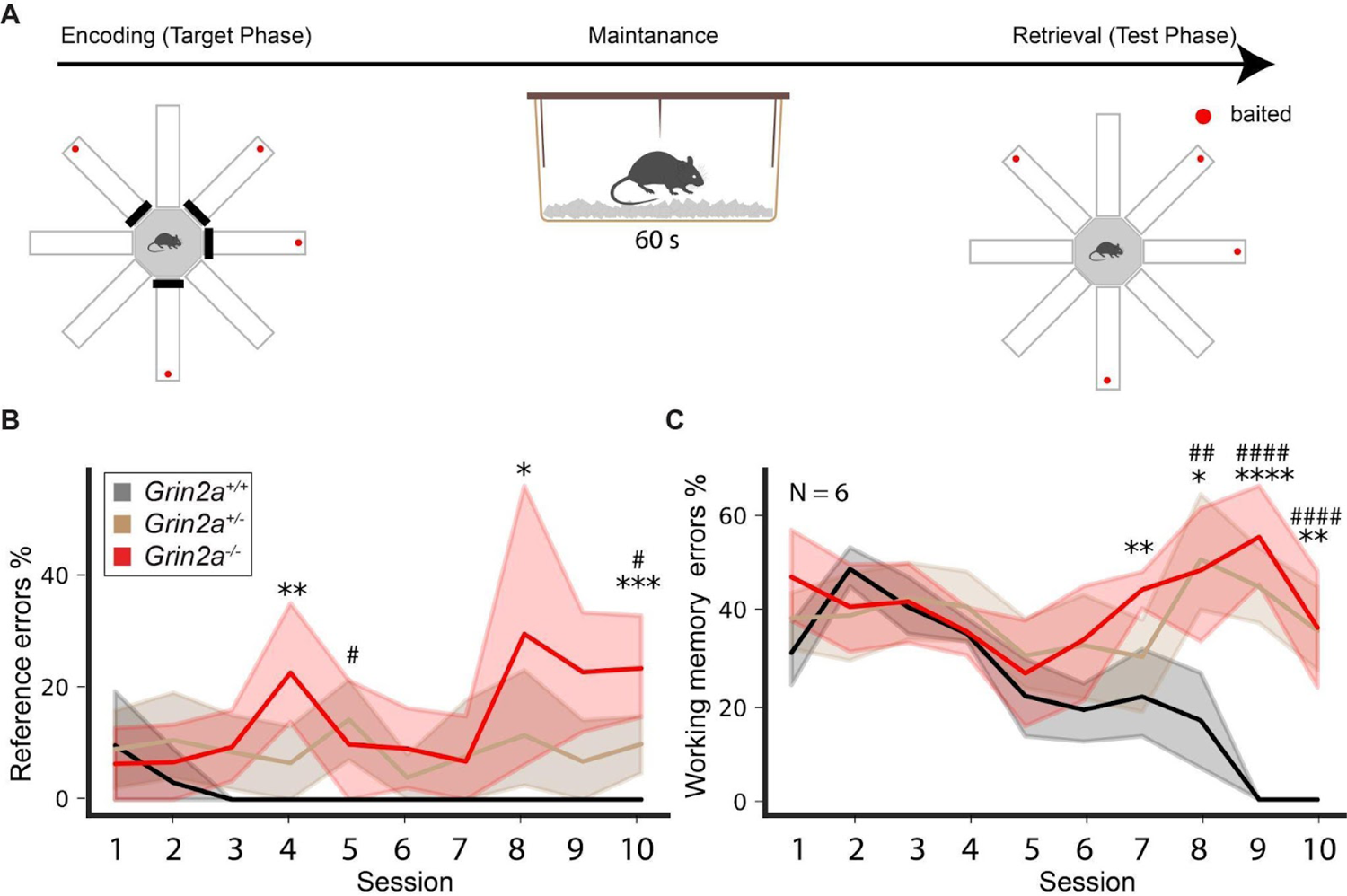
Spatial working memory is impaired in *Grin2a* mutants. **(A)** Schematic of the 8-arm radial maze paradigm used to test spatial working memory in mice. During the encoding phase, baited arms are pseudo-randomly blocked, followed by a 60-second maintenance period, and then the retrieval (test) phase trials. **(B)** Percentage of reference memory errors (incorrectly entering an arm that was never baited) across ten training sessions. Statistically significant differences in reference errors between wild-type (WT) and *Grin2a* mutants were observed during sessions S4 (WT: 0.0 ± 0.0, *Grin2a*⁺/⁻: 6.6 ± 4.1, *Grin2a*⁻/⁻: 22.8 ± 7.1), S5 (WT: 0.0 ± 0.0, *Grin2a*⁺/⁻: 14.4 ± 4.9, *Grin2a*⁻/⁻: 9.1 ± 5.0), S8 (WT: 0.0 ± 0.0, *Grin2a*⁺/⁻: 11.5 ± 6.8, *Grin2a*⁻/⁻: 29.7 ± 16.9), and S10 (WT: 0.0 ± 0.0, *Grin2a*⁺/⁻: 9.9 ± 3.5, *Grin2a*⁻/⁻: 23.6 ± 6.5). **(C)** Percentage of working memory errors (re-entering a previously entered arm within the same trial) across ten training sessions. Statistically significant differences in working memory errors between wild-type (WT) and *Grin2a* mutants were observed during sessions S7 (WT: 21.6 ± 6.5, *Grin2a*⁺/⁻: 30.0 ± 6.2, *Grin2a*⁻/⁻: 44.1 ± 2.4), S8 (WT: 16.6 ± 6.1, *Grin2a*⁺/⁻: 50.3 ± 7.9, *Grin2a*⁻/⁻: 48.2 ± 9.1), S9 (WT: 0.0 ± 0.0, *Grin2a*⁺/⁻: 44.7 ± 5.2, *Grin2a*⁻/⁻: 55.2 ± 7.3), and S10 (WT: 0.0 ± 0.0, *Grin2a*⁺/⁻: 35.5 ± 5.5, *Grin2a*⁻/⁻: 36.1 ± 8.5). Data are presented as mean ± SEM, with n = 6 mice per genotype. Statistical significance was assessed using independent t-tests (\**p* < 0.05, \*\**p* < 0.01, \*\*\**p < 0.001, ****p < 0.0001). Asterisks (\**) indicate comparisons between WT and *Grin2a*⁻/⁻, and hash symbols (#) indicate comparisons between WT and *Grin2a*⁺/⁻.

### Oscillations are Disrupted in Hippocampal Slices from *Grin2a* Mice

Radial maze performance requires hippocampal function both in rodents and humans (Goodrich-Hunsaker and Hopkins 2010). During maze tasks, hippocampal oscillations in the gamma and theta frequency bands increase and correlate with improved performance (Vugt et al. 2010; Yamamoto et al. 2014). We hypothesized that working memory deficits in *Grin2a* mutant mice may be linked to disruptions in hippocampal oscillations. oscillations. To test this, we conducted electrophysiological recordings on ex vivo hippocampal slices from WT and Grin2a mutant mice. Slices were placed on a multi-electrode array (MEA) chip with electrodes positioned in CA1 and CA3 to capture activity in these subfields (**Fig. 2A**). Under basal conditions, electrical activity in *ex vivo* hippocampal slices is low (Hermann and van Amsterdam 2015). Bath application of kainate increases activity in hippocampal sections, particularly in CA3 (Robinson and Deadwyler 1981; Castillo, Malenka, and Nicoll 1997). Kainate increased electrical activity throughout hippocampal sections from both WT and *Grin2a* mouse (**Fig. 2B, 2C, 2E**), but significantly increased gamma band oscillations (GBO) power in the CA3 region of *Grin2a-/-* (**Fig. 2E, 2F**). GBOs can dynamically couple within and between hippocampal subfields, with coupling strength increasing as cognitive load intensifies (Montgomery and Buzsáki 2007). We evaluated GBO coupling dynamics in WT and *Grin2a* slices, but did not detect any differences (**Fig. S2A-C**).

**Figure 2.**
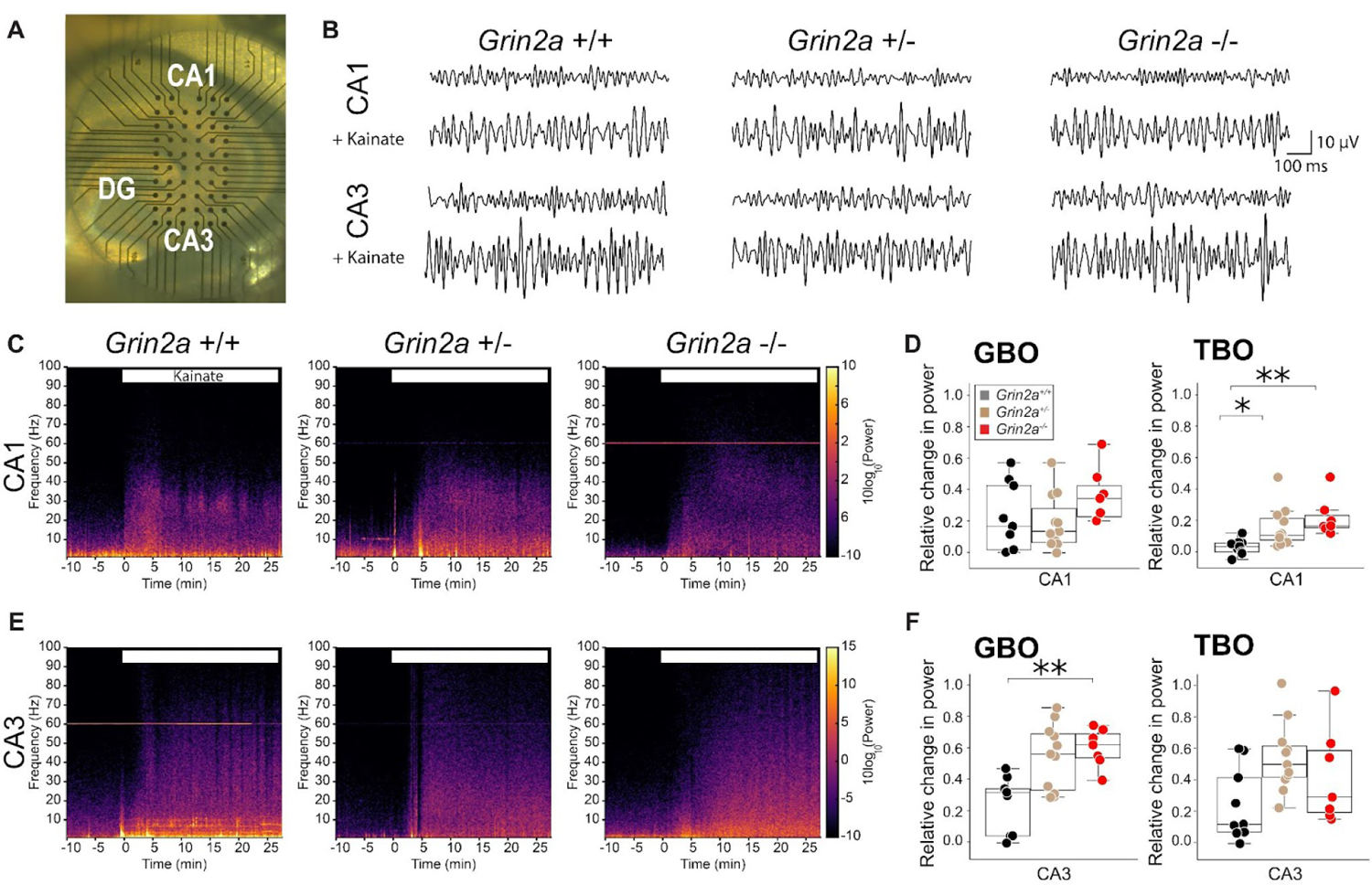
Hippocampal Oscillations are impaired in *Grin2a* mutants (A) Brightfield micrograph of an *ex vivo* hippocampal section mounted on a pMEA chip showing placement of electrodes in the CA1 and CA3 subfields. (B) LFP recordings from CA1 or CA3 electrodes. Top traces show baseline recordings; bottom panels show recordings during bath application of kainate. (C) Prototypical spectrograms of CA1 broadband changes in activity following kainate application for WT (*Grin2a*⁺/+, left), heterozygous (*Grin2a*⁺/⁻, middle), and knockout (*Grin2a*⁻/⁻, right) mice. (D) Kainate-induced changes in GBO and TBO power in CA1. GBO power values are WT (*Grin2a*+/+): 0.17 ± 0.02, *Grin2a*+/-: 0.13 ± 0.008, *Grin2a*-/–: 0.34 ± 0.01. For TBO power, values are WT (*Grin2a*+/+): −0.03 ± 0.003, *Grin2a*+/-: 0.1 ± 0.006 (p < 0.05), *Grin2a*-/–: 0.16 ± 0.007 (p < 0.01). Statistical significance was determined using the Kruskal-Wallis test, with p < 0.05 (*), p < 0.01 (**). Data are presented as mean ± SEM from 7-11 hippocampal slices per condition. (E) Prototypical spectrograms of CA3 broadband changes in activity following kainate application for WT (*Grin2a*⁺/+, left), heterozygous (*Grin2a*⁺/⁻, middle), and knockout (*Grin2a*⁻/⁻, right) mice. (F) Kainate-induced changes in GBO and TBO power in CA3. GBO power: WT (*Grin2a*+/+); 0.32 ± 0.01, *Grin2a*+/- 0.56 ± 0.01; *Grin2a*-/– 0.62 ± 0.006 (p < 0.001). TBO power: WT (*Grin2a*+/+) 0.25 ± 0.01; *Grin2a*+/- 0.5 ± 0.01; *Grin2a*-/– 0.43 ± 0.01. Statistical significance was determined using the Kruskal-Wallis test, with p < 0.05 (*), p < 0.01 (**). Data are presented as mean ± SEM from 7-11 hippocampal slices per condition.

Theta band oscillations (TBOs) are rhythmic oscillations within the 7-9 Hz frequency range. In the CA1 region, TBO power is inversely related to reference memory errors during radial maze tasks (Masuoka, Fujii, and Kamei 2006). Because Grin2a mice committed more reference memory errors than WT mice, we further characterized kainate-induced changes in TBO power. Kainate increased TBO power in the CA3 region of hippocampal sections across all genotypes (**Fig. 2E-F**) (Boehlen et al., 2009).

However, in CA1, kainate-induced genotype-specific effects on TBO power. In Grin2a+/- and Grin2a-/- mice, kainate increased CA1 TBO power by 35% and 37%, respectively, but reduced it by 34% in WT mice (p < 0.05 and p < 0.01) (**Fig. 2D**). Together, these results suggest that although the mechanisms of kainate-induced GBOs and their coupling dynamics remain largely intact in Grin2a hippocampal sections, TBO dynamics in CA1 exhibit genotype-specific alterations.

### Theta-Gamma Phase Amplitude Coupling is Attenuated in *Grin2a* Mutants

Theta-gamma phase-amplitude coupling (PAC) strength has been shown to correlate with working memory performance in previous studies (Tort et al. 2009; Tamura et al. 2017). GBOs are coupled to the phase of theta oscillations, with GBO amplitude peaking near the crest of the theta cycle (**Fig 3A**). PAC is preserved in *ex vivo* preparations (Colgin et al. 2009), and GluN2A subunit mutations are known to reduce the efficacy of this coupling (Bertocchi et al. 2021). To investigate the effects of Grin2a ablation on theta-gamma PAC, we generated spectrograms from hippocampal slices before and after kainate application. We quantified PAC strength using normalized direct PAC (ndPAC) in hippocampal slices from both WT and Grin2a mutant mice. Under vehicle conditions, theta-gamma PAC was significantly reduced in the CA1 and CA3 regions of Grin2a mutants compared to WT mice (**Fig. 3B-E**). Following kainate application, theta-gamma PAC significantly increased in the CA3 region of *Grin2a* mutants, while it remained unchanged in WT slices (**Fig. 3D, E**). Notably, kainate also induced a phase shift in PAC in both the CA1 and CA3 regions of *Grin2a* mutants, an effect not observed in WT mice (**Fig. S3**).

**Figure 3.**
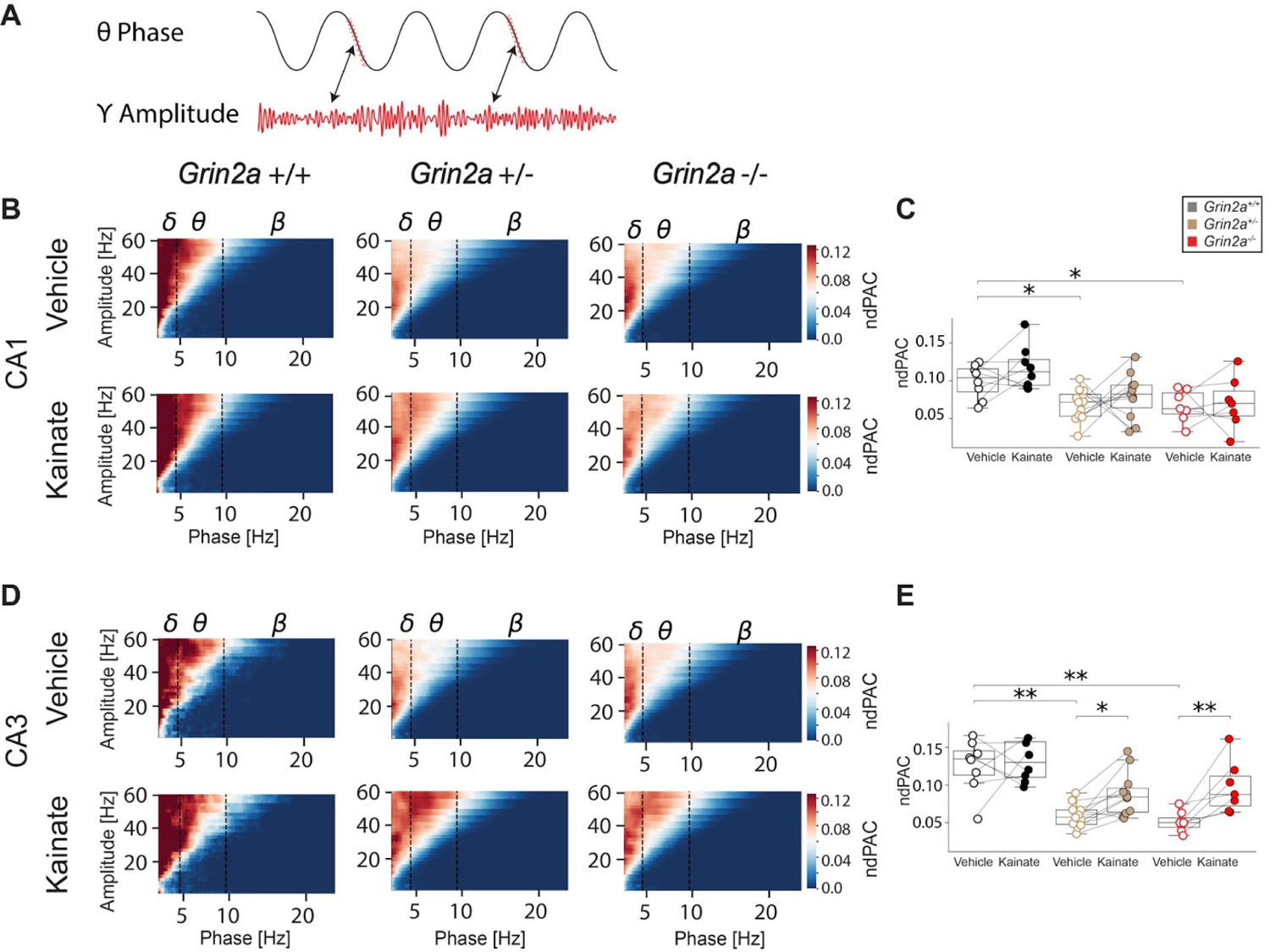
Kainate increases phase-amplitude coupling in the CA3 of *Grin2a* mutants **(A)** Schematic representation of theta-gamma PAC **(B)** Spectrograms of theta-gamma phase-amplitude coupling in CA1 hippocampal sections from *Grin2a* mutants in vehicle or kainate **(C)** Quantitative analysis of normalized direct slow gamma power relative to theta phase (ndPAC) in CA1. ndPAC (vehicle): *Grin2a*+/+: 0.1 ± 0.008, *Grin2a*⁺/⁻: 0.07 ± 0.007, *Grin2a*⁻/⁻: 0.06 ± 0.008. ndPAC (kainate): *Grin2a*+/+: 0.12 ± 0.01, *Grin2a*⁺/⁻: 0.08 ± 0.009, *Grin2a*⁻/⁻: 0.07 ± 0.01. **(D)** Spectrograms of theta-gamma phase-amplitude coupling in CA3 hippocampal sections from *Grin2a* mutants in vehicle or kainate **(E)** Quantitative analysis of normalized direct slow gamma power relative to theta phase (ndPAC) in CA3. ndPAC (vehicle): *Grin2a*+/+: 0.13 ± 0.01, *Grin2a*+/-: 0.06 ± 0.004, *Grin2a*-/-: 0.05 ± 0.005. ndPAC (kainate): *Grin2a*+/+: 0.14 ± 0.02, *Grin2a*+/-: 0.09 ± 0.008, *Grin2a*-/-: 0.1 ± 0.01. Data represent mean ± SEM from 7-11 hippocampal sections per genotype. Statistical significance was determined using the Kruskal-Wallis test, with p < 0.05 (*), p < 0.01 (**)

### Spike Coupling to Theta Phase is Impaired in *Grin2a* Mutant Mice

The timing of neuronal spikes relative to theta oscillation phase is crucial for hippocampal functions such as spatial encoding and memory (Harris et al. 2003; O’Keefe and Burgess 2005). To assess the impact of *Grin2a* deficiency on spike-phase coupling, we analyzed spike timing synchronization to theta oscillations in putative excitatory (Ep) and inhibitory (Ip) neuronal units. We classified Ep and Ip units using principal component analysis (PCA)-mediated spike sorting, combined with the amplitude of the after-hyperpolarization potential (**Fig 4A**). We quantified spike synchronization with theta oscillations using the phase locking value (PLV), a measure of how consistently spikes occur at specific theta phases (**Fig. 4B**). PLV values for both Ep and Ip units were significantly lower in *Grin2a* mutants than WT mice (**Fig. 4C-F**). This reduced synchronization suggests that *Grin2a* dysfunction disrupts the precise timing of neuronal spikes relative to local field potentials, potentially contributing to the observed deficits in hippocampal-dependent functions.

**Figure 4.**
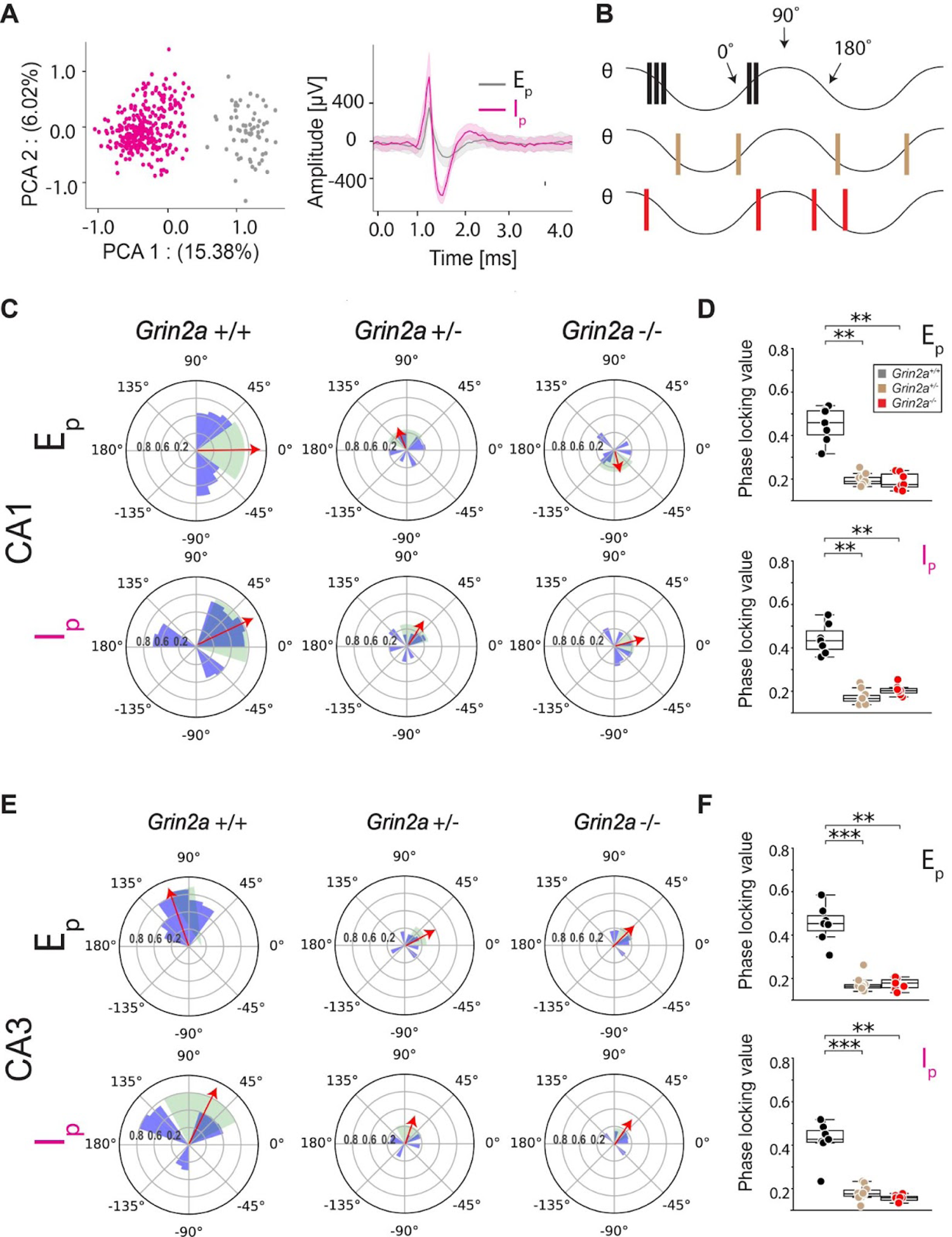
Spike-phase coupling is attenuated in the hippocampus of *Grin2a* mice **(A)** Principal component analysis (PCA) and spike sorting were utilized to categorize LFP waveforms into putative excitatory units (E_p_) and putative inhibitory units (I_p_). **(B)** Schematic representation of spike coupling to the theta phase of the LFP, WT (black), heterozygous (tan), or homozygous *Grin2a* mice (red). **(C)** Angular histogram displaying spike coupling to the theta phase of E_p_ or I_p_ units in the CA1 of *Grin2a*+/+, *Grin2a*+/-, and *Grin2a*-/- mice. Red arrows indicate the mean phase locking value (PLV); green shading represents the standard deviation of PLV. *Note the difference in scales used for WT and Grin2a mutants*. **(D)** Box plots representing phase-locking values (PLV) for CA1 putative excitatory (E_p_) and putative inhibitory (I_p_) units by genotype. For E_p_ units values were *Grin2a*+/+: 0.38 ± 0.04; *Grin2a+/-*: 0.2 ± 0.02; and *Grin2a-/-*: 0.2 ± 0.02. For I_p_ units values were *Grin2a*+/+: 0.39 ± 0.05, *Grin2a+/-*: 0.16 ± 0.01; *Grin2a-/-*: 0.16 ± 0.01. Data are presented as mean ± SEM, with 8–9 hippocampal slices per genotype. Statistical significance was determined using the Kruskal-Wallis test, with p < 0.05 (*), p < 0.01 (**), p < 0.001 (***). **(E)** Angular histogram displaying spike coupling to the theta phase of E_p_ or I_p_ units in the CA3 of *Grin2a*+/+, *Grin2a*+/-, and *Grin2a*-/- mice. Red arrows indicate the mean phase locking value (PLV); green shading represents the standard deviation of PLV. *Note the difference in scales used for WT and Grin2a mutants*. **(F)** Box plots representing phase-locking values (PLV) for CA3 putative excitatory (E_p_) and putative inhibitory (I_p_) units by genotype. For E_p_ units values were *Grin2a*+/+: 0.45 ± 0.03; *Grin2a+/-*: 0.20 ± 0.01; and *Grin2a-/-*: 0.19 ± 0.01. For I_p_ units values were *Grin2a*+/+: 0.44 ± 0.03, *Grin2a+/-*: 0.17 ± 0.01; *Grin2a-/-*: 0.20 ± 0.01. Data are presented as mean ± SEM, with 7–11 hippocampal slices per genotype. Statistical significance was determined using the Kruskal-Wallis test, with p < 0.05 (*), p < 0.01 (**), p < 0.001 (***).

### PV+ Neuron Density and Inhibitory Tone are increased in the CA1 region of *Grin2a* Mice

PV+ interneurons are essential for regulating spike-phase coupling to theta oscillations in the hippocampus, a process critical for cognitive function (Strüber, Sauer, and Bartos 2022), which is essential for cognitive function (Sohal et al 2012).Previous studies indicate that Grin2a ablation increases PV+ neuron density specifically in the CA1 region (Camp et al. 2023b). To confirm the reported increase in PV+ neuron density and assess its impact on inhibitory tone, we used immunohistochemistry (IHC) to quantify PV+ cell abundance in CA1 and CA3, and electrophysiology to directly measure inhibitory tone in CA1. IHC confirmed increased PV+ neuron abundance in the CA1 region and showed preserved hippocampal morphology in *Grin2a* mutants (**Fig. 5A**).

**Figure 5.**
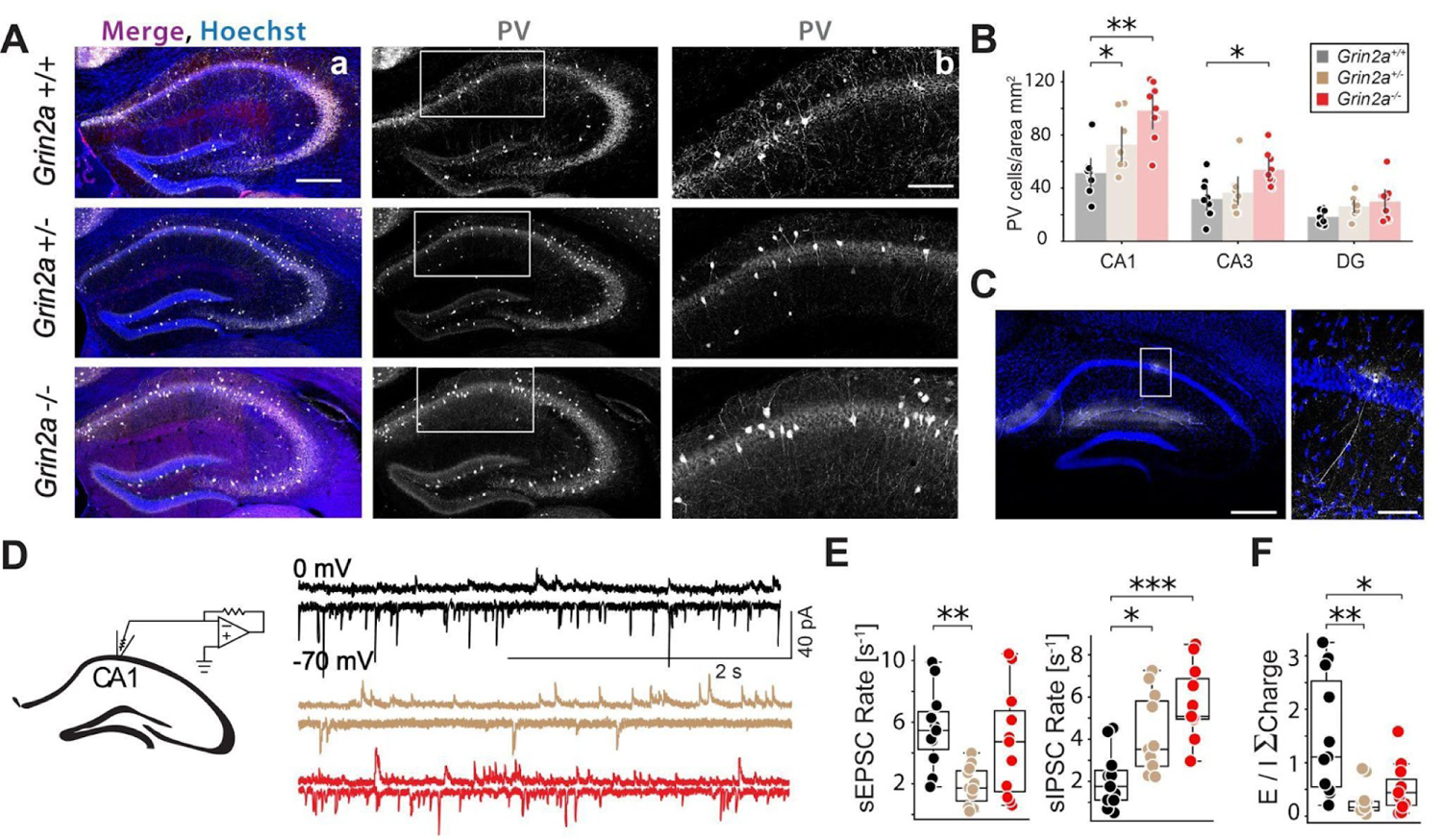
PV+ staining and inhibitory tone is increased in the CA1 of *Grin2a* mutants. **(A)** Representative confocal micrographs of hippocampal sections immunostained for parvalbumin (PV) (white) and Hoechst (blue) across *Grin2a*-/-, *Grin2a*+/-, and *Grin2a*+/+ mice. Scale bars: 200 μm for (a) columns, and 100 μm for (b) columns. **(B)** Quantitative analysis of PV-positive cells in CA1 and CA3 *Grin2a*-/- mice compared to *Grin2a*+/+ and *Grin2a*+/- mice. N = 3 mice per genotype, *n* = 3-4 sections per brain. CA1: WT (*Grin2a*+/+): 51.3 ± 5.5, *Grin2a*+/-: 72.7 ± 6.6, *Grin2a*-/-: 98.3 ± 7.1. For CA3: WT *Grin2a*+/+: 31.8 ± 4.8, *Grin2a*+/-: 36.6 ± 5.5, *Grin2a*-/-: 53.8 ± 4.3. For DG: WT *Grin2a*+/+: 18.3 ± 1.8, *Grin2a*+/-: 26.3 ± 2.5, *Grin2a*-/-: 29.7 ± 4.7. Data presented as mean ± SEM. Statistical significance denoted as *p < 0.05, **p < 0.01, using the Kruskal-Wallis Test. **(C)** Confocal images of a biocytin-filled pyramidal neuron in layer V of WT CA1. **(D)** Schematic illustration of whole-cell patch clamp recording from a CA1 pyramidal cell; representative traces showing spontaneous excitatory (sEPSC at −70mV) and inhibitory postsynaptic currents (sIPSC at 0mV) in CA1 pyramidal cells. **(E)** Box plots of mean frequency (in Hz) of sEPSC: WT *Grin2a*+/+: 5.57 ± 0.76, *Grin2a*+/-: 1.88 ± 0.38; *Grin2a*-/-: 4.7 ± 1.06 and sIPSC: WT *Grin2a*+/+: 2.07 ± 0.4, *Grin2a*+/-: 4.18 ± 0.56; *Grin2a*-/-: 5.73 ± 0.53 rates in CA1. **(F)** Box plots sEPSC/sIPSC ratio of of total charge transfer: WT *Grin2a*+/+: 1.5 ± 0.33, *Grin2a*+/-: 0.29 ± 0.09; *Grin2a*-/-: .53 ± 0.13 in CA1. Data represent measurements from 11-14 cells from three mice. Statistical significance is denoted by asterisks: *p < 0.05, **p < 0.01, ***p < 0.001, determined via the Kruskal-Wallis test.

These histological findings were complemented by electrophysiological measurements that directly assessed changes in inhibitory tone linked to Grin2a dysfunction. Quantification of PV+ cells showed a ∼41.7% increase in *Grin2a*+/- and a ∼91.6% increase in *Grin2a*-/- mutants compared to WT in the CA1 region (**Fig. 5B**). The CA3 region of *Grin2a*-/- mutants also exhibited a ∼69.2% increase in PV+ cell density relative to WT (**Fig. 5B**).

Given the altered network activity in G*rin2a* mice, we next conducted voltage-clamp recordings to measure spontaneous excitatory (sEPSCs) and inhibitory (sIPSCs) postsynaptic currents in CA1 pyramidal neurons (**Fig. 5C, D**). The frequency of sIPSCs was significantly higher in both *Grin2a*+/- and *Grin2a*-/- mice (**Fig. 5E**), though no changes were detected in amplitude or decay time (**Fig. S4B**). These results suggest that the increased PV+ cells integrate successfully into hippocampal circuits and form functional inhibitory synapses. In contrast, the frequency of sEPSCs in CA1 pyramidal neurons was lower in *Grin2a*+/- mice compared to WT (**Fig. 5E**), and sEPSC amplitude was reduced in both *Grin2a*+/- and *Grin2a*-/- mice (**Fig. S4A**). Additionally, sEPSC decay times were prolonged in *Grin2a* mutants (**Fig. S4A**), consistent with previous reports (Booker et al. 2021).

As both excitatory (e.g., prolonged decay times) and inhibitory changes (e.g., decreased frequency and amplitude of sEPSCs) were observed across genotypes, the cumulative impact of *Grin2a* dysfunction on net synaptic activity remained unclear. To address this, we developed a computational model integrating variations in sEPSC amplitude, frequency, and kinetics, providing a quantitative approach to evaluate the overall effect on charge transfer. Despite these excitatory changes, our model indicates that *Grin2a* dysfunction leads to a net increase in inhibitory charge transfer in CA1 pyramidal neurons (**Fig. 5F**).

## Discussion

This study demonstrates that both complete and partial loss of GluN2A-containing NMDARs significantly impair hippocampal-dependent working memory, disrupt network oscillations, and alter excitatory-inhibitory (E/I) signaling. *Grin2a* mutant mice, whether lacking one or both gene copies, showed persistent deficits in cognitive flexibility and working memory. We identified disruptions in theta-gamma PAC and PV+ neuron dynamics as critical contributors to these impairments. Notably, our findings reveal a non-linear gene-dose response, with *Grin2a* heterozygous mutants showing cognitive and physiological deficits comparable to homozygous mutants, emphasizing the significant impact of even partial GluN2A dysfunction on network stability.

*Grin2a* mutant mice exhibited deficits in both working memory and cognitive flexibility. Persistent reference errors—entries into unbaited arms—indicate impaired cognitive flexibility, as the task required adaptive strategies in response to changing bait configurations. By the third session, WT mice made significantly fewer reference errors, reflecting successful task adaptation, while *Grin2a* mutants continued making errors throughout. These impairments align with previous findings linking GluN2A dysfunction to deficits in set-shifting and discrimination reversal (Marquardt et al. 2014).

Our findings suggest that spatial working memory deficits in *Grin2a* mutants are linked to disruptions in hippocampal network oscillations, particularly reductions in GBO power, spike-phase coupling, and theta-gamma phase-amplitude coupling (PAC). These oscillatory dynamics are crucial for synchronizing neuronal activity during cognitive tasks (Buzsáki and Wang 2012). Disruptions in theta-gamma PAC impair neuronal ensemble coordination, which may contribute to the observed deficits in cognitive flexibility and working memory (Lisman and Jensen 2013). Such oscillatory disruptions likely hinder *Grin2a* mutants from forming adaptive search strategies.

Our findings indicate that kainate has region- and genotype-specific effects on GBO and TBO power in the hippocampus. Kainate application increased GBO power in both WT and *Grin2a* mutant slices, with a notably stronger effect in the CA3 region of *Grin2a* mutants, where GBO power nearly doubled relative to WT. The heightened CA3 sensitivity observed in *Grin2a* mutants is consistent with prior studies showing CA3’s increased responsiveness to kainate(Castillo, Malenka, and Nicoll 1997).

Theta-gamma phase-amplitude coupling (PAC), essential for hippocampal-dependent functions like spatial working memory (Tort et al. 2009), was reduced in *Grin2a* mutants under basal conditions. This reduction may result from an imbalance in the excitatory-inhibitory (E/I) ratio, as increased inhibitory input from PV+ GABAergic interneurons weakens cross-frequency coupling and disrupts theta-gamma synchrony (Sotero 2016). Such heightened inhibitory tone in *Grin2a* mutants appears to impair hippocampal network coherence and may contribute to cognitive deficits.

The heightened sensitivity of CA3 to kainate in *Grin2a* mutants likely amplifies kainate’s effects on GBO and TBO power in both CA3 and CA1. As a primary driver of GBOs in CA1 (Colgin et al. 2009), CA3’s more robust response to kainate in *Grin2a* mutants may lead to increased input to CA1, contributing to the elevated GBO and TBO power observed there. In WT mice, kainate did not alter TBO power in either CA3 or CA1, highlighting a genotype-specific impact of *Grin2a* dysfunction on hippocampal network dynamics.

Consistent with previous studies (Camp et al. 2023), our findings reveal an increase in PV+ interneuron density throughout the hippocampus of *Grin2a* mice, particularly in CA1 and CA3. This heightened inhibitory input to CA1 pyramidal neurons disrupts the excitatory-inhibitory (E/I) balance, impairing oscillatory coherence and weakening long-range network synchronization. These disruptions, evidenced by reduced GBO coherence and attenuated theta-gamma PAC, likely interfere with the precise timing of neuronal activity needed for spatial and working memory, contributing to the cognitive deficits observed in *Grin2a* mice.

Cognitive impairments in *Grin2a* heterozygous mice are less pronounced than those observed in homozygous knockouts, though both genotypes exhibit highly similar transcriptomic changes in excitatory neurons in CA1 and CA3 (Farsi et al. 2023). This pattern suggests that even partial loss of GluN2A leads to extensive molecular changes, contributing to network disruptions, including impairments in GBO power and theta-gamma PAC. These findings align with previous reports of EEG alterations in both heterozygous and homozygous *Grin2a* mice (Herzog et al. 2023). The increased network instability observed in *Grin2a* mutants may also heighten the risk of epileptiform activity, consistent with associations between GluN2A mutations and epilepsy-aphasia spectrum disorders (Bertocchi et al. 2021). These results underscore the translational relevance of the *Grin2a* model, as human patients with a single GRIN2A allele mutation display comparable cognitive and network deficits.

While *ex vivo* hippocampal slices reveal fundamental cellular mechanisms, they cannot fully capture the complex *in vivo* interactions between the hippocampus and other brain regions, such as the prefrontal cortex, crucial for cognitive function. Future *in vivo* studies using techniques like electrophysiology, optogenetics, or fMRI will be needed to track hippocampal-prefrontal connectivity in real-time, particularly for understanding the role of long-range network synchrony in cognition. Additionally, reliance on a single genetic mouse model may not represent the full spectrum of GluN2A mutations seen in humans (Elmasri et al. 2022). Investigating a more comprehensive range of mutations, including missense variants associated with epilepsy-aphasia spectrum disorders, will be essential for understanding how GluN2A dysfunction contributes to diverse neuropsychiatric and neurological conditions (Bertocchi et al. 2021).

This study underscores the critical role of GluN2A-containing NMDARs in regulating hippocampal network dynamics and cognitive function. Disruptions in GBO coherence, increased PV+ interneuron density, and attenuated theta-gamma PAC illustrates how imbalances in excitatory-inhibitory signaling may underlie deficits in working memory and cognitive flexibility. These findings offer mechanistic insights into the neural basis of conditions like schizophrenia and epilepsy, where network synchronization is disrupted. Therapeutic strategies to restore network synchrony hold promise for cognitive improvement. Testing NMDAR modulators, such as D-cycloserine or PV+ interneuron-targeted therapies in preclinical models could help determine if restoring oscillatory coherence improves working memory and cognitive flexibility while reducing seizure susceptibility, potentially addressing cognitive impairments in disorders linked to GluN2A dysfunction.

K.S.J. conceptualized and designed the study. H.H.,and J.C.R..D conducted the study and performed analysis. K.S.J secured the funding. H.H.,and J.C.R..D developed the methodology and carried out the experiments. H.H constructed apparatuses. K.S.J and D.L.P oversaw project administration and supervision. H.H and J.C.R.D were responsible for the visualization. J.C.R.D., H.H. and K.S.J wrote the original draft of the manuscript. All authors reviewed and edited the manuscript, contributing to the interpretation of the results and providing critical feedback throughout the research process.This work was supported by

---

**Figure S1.**
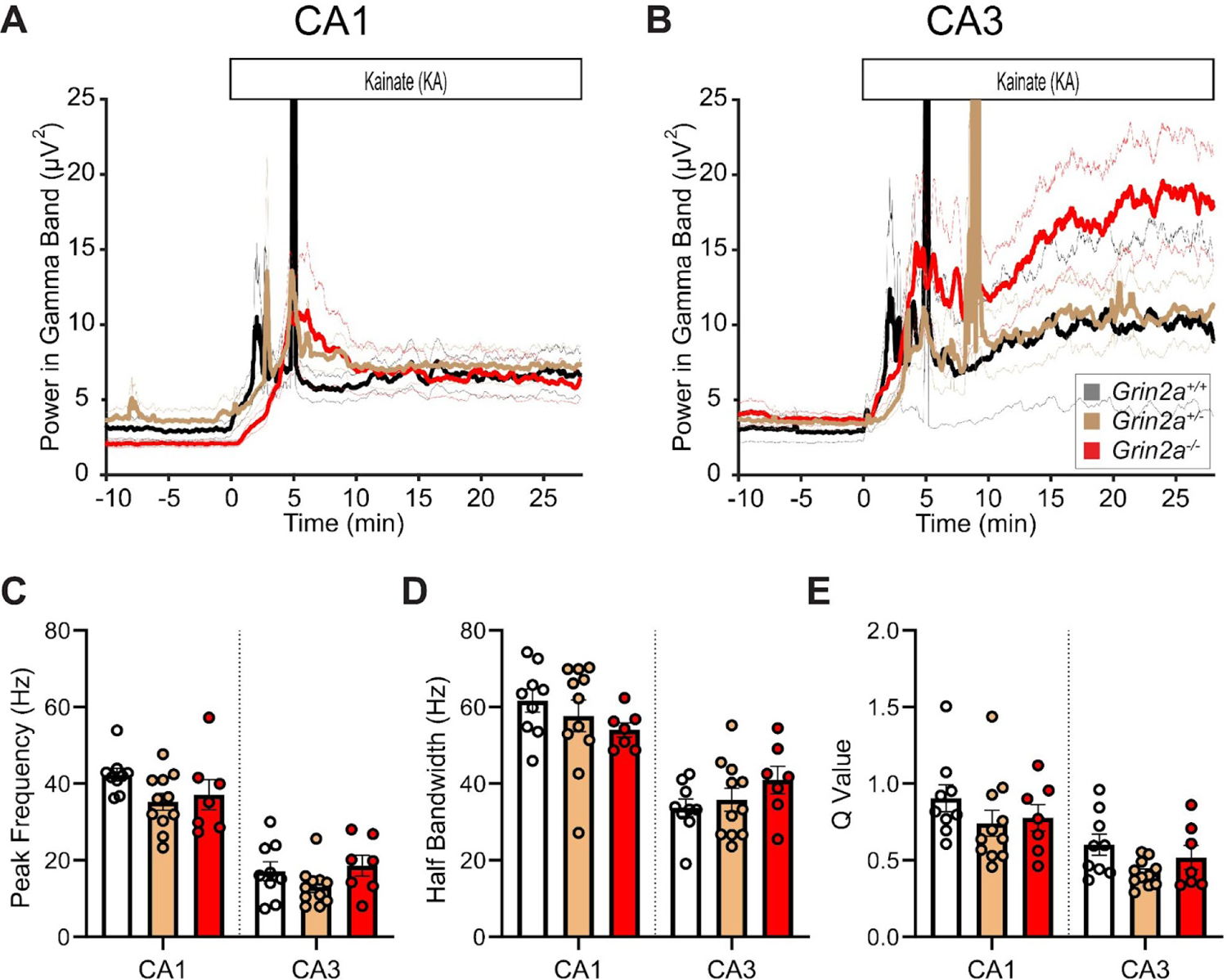
Kainate-evoked GBOs are stronger in *Grin2a* KO mice. **(A, B)** Time course of kainate-induced changes in GBO power in CA1 and CA3 of WT and *Grin2a* mutants. *Grin2a +/+* (black), *Grin2a*+/- (beige), and *Grin2a* -/- (red) Data are represented as mean ± SEM, where *n* = number of slices. Statistical significance denoted by asterisks: *p < 0.05, **p < 0.01, ***p < 0.001, ****p < 0.0001. **(C)** Peak frequency of kainate-evoked oscillations in CA1 and CA3: **CA1**: *Grin2a*+/+ (WT): 42.3 Hz ± 1.70 (*n* = 9), *Grin2a*+/-: 35.3 Hz ± 2.23 (*n* = 11), *Grin2a*-/-: 37.1 Hz ± 4.0 (*n* = 7), ANOVA (F = 2.16, p = 0.14); CA3: *Grin2a*+/+: 17.2 Hz ± 2.47 (*n* = 9), *Grin2a*+/-: 13.2 Hz ± 1.51 (*n* = 11), *Grin2a*-/-: 18.6 Hz ± 2.76 (*n* = 7), ANOVA (F = 1.79, p = 0.19). **(D)** Half bandwidth of GBOs in CA1 and CA3. **CA1**: *Grin2a*+/+ (WT): 61.7 Hz ± 3.05 (*n* = 9), *Grin2a*+/-: 57.7 Hz ± 4.12 (*n* = 11), *Grin2a*-/-: 54.0 Hz ± 1.88 (*n* = 7), ANOVA (F = 1.04, p = 0.37); CA3: *Grin2a*+/+: 33.7 Hz ± 2.30 (*n* = 9), *Grin2a*+/-: 35.8 Hz ± 3.01 (*n* = 11), *Grin2a*-/-: 41.0 Hz ± 3.58 (*n* = 7), ANOVA (F = 1.35, p = 0.28). **(E)** Quality (Q) values of oscillations in CA1 and CA3 for *Grin2a*+/+, *Grin2a*+/-, and *Grin2a*-/- genotypes. **CA1**: *Grin2a*+/+ (WT): 0.90 ± 0.09 (*n* = 9), *Grin2a*+/-: 0.74 ± 0.09 (*n* = 11), *Grin2a*-/-: 0.78 ± 0.09 (*n* = 7), ANOVA (F = 0.98, p = 0.39); CA3: *Grin2a*+/+: 0.60 ± 0.07 (*n* = 11), *Grin2a*+/-: 0.42 ± 0.03 (*n* = 11), *Grin2a*-/-: 0.52 ± 0.08 (*n* = 7), ANOVA (F = 3.09, p = 0.06).

**Figure S2.**
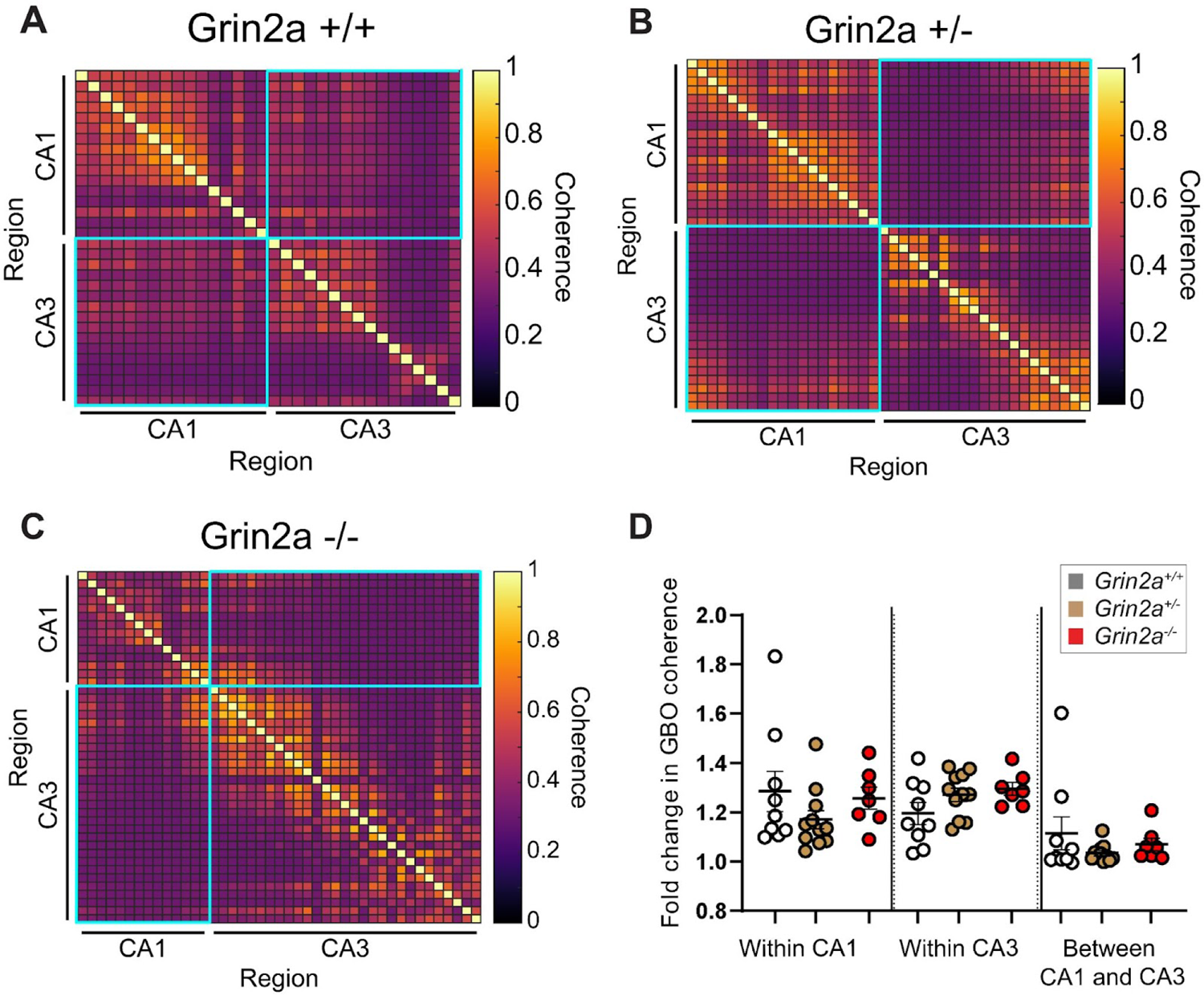
Spatial organization of GBO power coherence in hippocampal sections from *Grin2a* mutants **(A-C)** Heat maps illustrating GBO power-power coherence in hippocampal sections from *Grin2a* genotypes: (A) *Grin2a*+/+, (**B**) *Grin2a*+/-, and (**C**) *Grin2a*-/-. Coherence between CA1 and CA3 regions is highlighted in cyan. **(D)** Quantitative analysis of the mean fold change in GBO coherence after GBO induction: within CA1 (+/+: 1.29 ± 0.08, n = 9; +/-: 1.17 ± 0.04, n = 11; -/-: 1.26 ± 0.04, n = 7), within CA3 (+/+: 1.20 ± 0.04, n = 9; +/-: 1.27 ± 0.03, n = 11; -/-: 1.30 ± 0.03, n = 7), and between CA1 and CA3 (+/+: 1.12 ± 0.07, n = 9; +/-: 1.03 ± 0.0, n = 11; -/-: 1.07 ± 0.03, n = 7). Statistical analyses via ANOVA indicated no significant genotype effects (F = 1.162.13, p = 0.31126) but significant regional differences (F = 16013.3164, p < 0.0001). Tukey’s multiple comparisons for each region are detailed.

**Figure S3.**
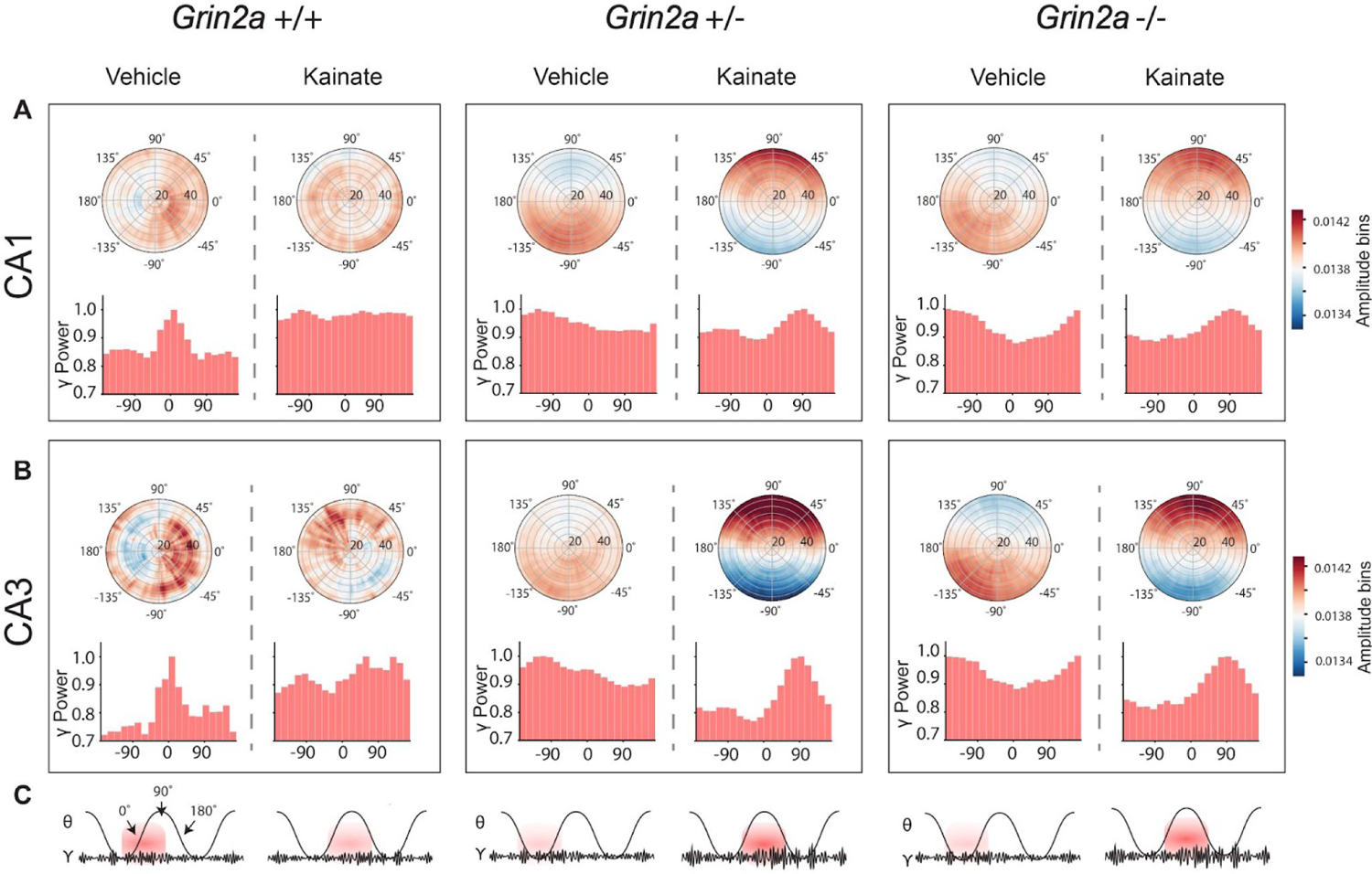
Kainate phase-shifts and strengthens theta-gamma in the CA1 and CA3 of *Grin2a* mutants **(A)** Polar plots displaying the distribution of baseline and kainate-induced slow gamma power relative to theta phase in the CA1 region for *Grin2a* -/-, *Grin2a* +/-, and *Grin2a* +/+ mice. Each plot comprises 72 bins representing 20-degree segments of the theta cycle (Top). Histograms showing the distribution of slow gamma power across theta phase bins, providing a quantitative analysis of how gamma power varies with theta phase under different genetic conditions (Bottom). **(B)** Polar plots displaying the distribution of baseline and kainate-induced slow gamma power relative to theta phase in the CA1 region for *Grin2a* -/-, *Grin2a* +/-, and *Grin2a* +/+ mice (Top). Histograms depicting the distribution of slow gamma power across theta phase bins, providing a quantitative analysis of gamma power variations within each theta cycle across different genotypes (Bottom). **(C)** Cartoon illustration of generalized changes in hippocampal theta-gamma PAC before and during kainate application by genotype

**Figure S4.**
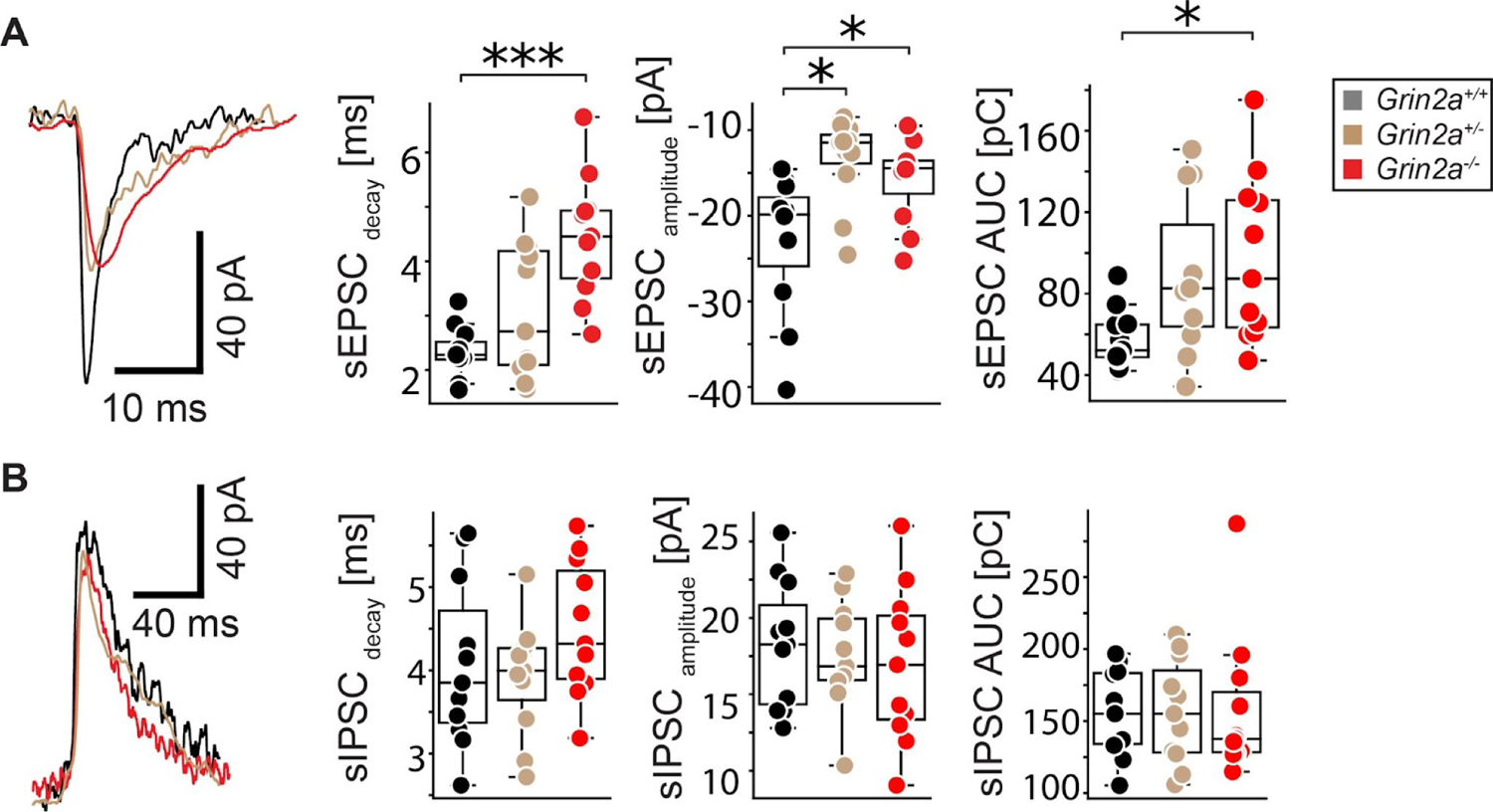
The amplitude and kinetics of sEPSCs and sIPSCs in CA1 *Grin2a* mutants. **(A)** Representative trace of a single sEPSC complemented by box plots depicting the kinetics, amplitudes and area under the curve (AUC) of sEPSCs in CA1 pyramidal cells across different *Grin2a* genotypes. **(B)** Representative trace of a single sIPSC complemented by box plots illustrating the kinetics, amplitudes and AUC of sIPSCs in CA1 pyramidal cells across different *Grin2a* genotypes. Data represent measurements from 11-14 cells derived from three mice per genotype. Statistical analysis was performed using the Kruskal-Wallis Test. Statistical significance denoted by asterisks: *p < 0.05, **p < 0.01, ***p < 0.001..

**Table 1.** Effect of Kainate on Oscillations in CA1 and CA3 by Genotype.

| Effect of Kainate on Oscillations in CA1 and CA3 by Genotype |  |  |  |  |  |
| --- | --- | --- | --- | --- | --- |
| Parameter | Region | Grin2a+/+ (WT) (9) | Grin2a+/- (11) | Grin2a-/- (11) | ANOVA |
| Baseline GBO Power ($\mu V^2$) | CA1 | 2.30 ± 0.14 | 3.02 ± 0.35 | 2.11 ± 0.24 | F = 2.97, p = 0.07 |
| | CA3 | 3.31 ± 0.58 | 3.46 ± 0.47 | 3.54 ± 0.43 | F = 0.046, p = 0.96 |
| Kainate-Evoked GBO Power ($\mu V^2$) | CA1 | 4.88 ± 0.41 | 6.52 ± 0.92 | 7.07 ± 1.33 | F = 1.45, p = 0.25 |
| | CA3 | 12.71 ± 4.83 | 10.66 ± 2.23 | 17.42 ± 3.58 | F = 0.858, p = 0.45 |

Data are presented as mean ± SEM, where (n) = number of slices

